# Implications of Endogenous Small Regulatory RNAs on Gene Silencing in Mollusks

**DOI:** 10.1101/2025.05.19.654968

**Authors:** Cory Von Eiff, Beatriz Schueng Zancanela, Megan Gima, Kevin Quito, Manitejus Kotikalapudi, Sergio Valdivia, Yulica Santos-Ortega, Alex Sutton Flynt

**Affiliations:** School of Biological Environmental and Earth Sciences, University of Southern Mississippi, 39406; Department of Biomedical Engineering, University of Mississippi, 38677; Thad Cochran Marine Aquaculture Center, Office of the Vice President for Research, University of Southern Mississippi, 39564; MD Anderson Cancer Biology Department, UT Health MD Anderson GSBS, 77030

**Keywords:** Mollusks, RNAi, sRNAs, piRNAs, Lophotrochozoa

## Abstract

Mollusks are an abundant group of animals with many economically important members that are phylogenetically distinct from nearly all genetic model organisms. This study provides clade-wide evaluation of sRNA biogenesis pathways, with emphasis on the easter oyster, *Crassostrea virginica*. Understanding these molecules prescribes RNAi-based gene silencing approaches, benefiting genetic investigation and biotechnology. Similar to other animal groups, mollusks have conserved microRNAs (miRNAs) with some shared with ecdysozoans and deuterostomes; however, there was no evidence of an endogenous small-interfering RNA (siRNA) pathway. These results suggest that long double-stranded RNA (dsRNA)-based RNAi is not appropriate for gene silencing in Mollusks as well as other members of the broader Lophotrochozoan clade. The study also finds an abundance of piwi-interacting RNAs (piRNAs) in both soma and gonads. Differences are also found in piRNA biology. Many invertebrates exhibit somatic piRNAs; however, mollusk piRNAs appear to be restricted to a subset of cells, limiting the potential of piRNA-based RNAi. Further, individual animals also express a unique collection of piRNAs that seem to be only partially determined through inheritance from both parents. Together this work defines the RNAi mechanisms in mollusks, which represent 23% of animals, and provides insights into the phenotypic diversity seen in this group.

**Significance Statement:** This study provides an extensive, clade-wide evaluation of small RNA (sRNA) biogenesis pathways in mollusks. Our findings reveal that, unlike ecdysozoans and deuterostomes, mollusks lack a functional siRNA pathway, which fundamentally changes expectations around RNA interference (RNAi) applications in Mollusca. Instead, we find the expected microRNAs and an assortment of piwi-interacting RNAs (piRNAs). We show that piRNA biology in mollusks is highly cell-type specific and genetically individualized. We further demonstrate that piRNA expression is likely linked to stem-like, quiescent cells, suggesting a critical role in genomic maintenance. This work offers insight into RNAi potential in mollusks, the second largest animal phylum, and has significant implications for both basic biology and applied sciences such as pest control and aquaculture.

## Introduction

Mollusks are an abundant and diverse group of animals with many economically important members that are the basis of major fisheries. An average of $20.6 billion dollars of economic activity from 2010-2015 was associated with mollusk fisheries, accounting for 14% of total marine fisheries (1). Unlike the other species, ∼90% of the bivalve harvest were from aquaculture. As there is extensive human interaction with animals, there is an opportunity to integrate biotechnologies into husbandry practices to enhance yields. Possibly the most facile genetic technology to deploy is RNA interference (RNAi) due to the simple nature of double-stranded RNAs typically used to trigger this process and elicit gene silencing. In animals, three RNAi pathways have been described: microRNAs (miRNAs), small interfering RNAs (siRNAs), and piwi-interacting RNAs (piRNAs) (2–4). Each pathway has been used for gene silencing through leveraging distinct biogenesis pathways and functions by guiding exogenous RNAs into respective effector Argonaute (Ago)/Piwi proteins (5–7). Differences in these pathways are often observed at the order level but have been seen between species (8, 9). Thus, successful RNAi benefits from understanding the endogenous small RNA biology of target species(9).

Among all major animal groups, sRNA pathways have been relatively unexplored in the Mollusca phylum and the broader Lophotrochozoan clade (10). Of these pathways, miRNAs are the most conserved, being ubiquitously identified in eukaryotes, including Mollusks (11). Originating from “short” hairpins, miRNAs are cropped by Drosha and cleaved by a miRNA processing Dicer (miDicer) (12–15). Both Drosha and Dicer are RNase III class enzymes that leave 2nt 3’ overhangs on small RNA duplex products (16, 17). After processing, miRNAs load into miRNA-specific Ago (miAgo) proteins and initiate degradation of mRNA by inhibiting translation or occasionally directing cleavage of the bound RNA, thus regulating gene expression (11, 18). In contrast, siRNAs, which are reported in many ecdysozoans, originate from long double-stranded RNA (dsRNA) and, after processing by a dedicated siRNA generating Dicer (siDicer), interact with siRNA-specific AGO (siAGO) proteins to initiate target cleavage. Often, siRNAs have anti-viral function, though there are many endogenous siRNAs (endo-siRNAs) that have cryptic roles that include genome maintenance and gene regulatory networks (7, 19–22). Endo-siRNAs are poorly conserved, being mostly absent in vertebrates and the annelid *Capitella teleta* (8, 15, 23).

The third class, piRNAs, are present in nearly all animals, including in Mollusks, with a notable exception of dust mites and some platyhelminths (10, 24, 25). Unlike miRNAs and siRNAs, piRNAs are not derived from dsRNA precursors (26, 27). Lack of Dicer processing and differences between the small RNA binding pocket of Ago and Piwi proteins leads to piRNAs having a distinct size of ∼26-30 nt (28, 29). piRNAs originate through two mechanisms: phasing and Ping Pong (30). Both pathways involve the interplay of Piwi partner proteins, which we define here as phasing Piwi (phPiwi) and responder Piwi (rePiwi) based on their respective role in phasing piRNA biogenesis. In phasing, a responder piRNA directs cleavage of a transcript that becomes a substrate for the Zucchini/mito-PLD (Zuc) RNase (27, 31, 32). Zuc processing yields a distinctive head to tail arrangement, thus the designation as “phasing” piRNAs. Cleavage usually occurs at uridine residues, leading to 1 U bias of phasing piRNAs. Using these features, phasing piRNAs can be recognized in small RNA sequencing alignments when reads show 1U and are arranged end to end (33–35). Ping Pong piRNAs are the result of an amplification loop coordinated by phPiwi and rePiwi. Transcripts cleaved by phPiwi become precursors of rePiwi-bound piRNAs and vice versa, thus leading to the loop (36, 37). piRNAs generated by Ping Pong have a characteristic 10 nt overlap, which can also be observed in read alignments (3). piRNAs have been found to silence transposable elements and mRNAs in a variety of animals (38–41). piRNAs are expressed in both somatic and gonadal tissue of Mollusks (10, 42).

Of the 75,000 extant Mollusk species, RNAi-mediated gene silencing has been reported in fewer than two dozen (43, 44). Most attempts involve soaking or injection of long synthetic dsRNAs, with the intention of these molecules to be processed into siRNAs and silence targets, as occurs in ecdysozoans like *D. melanogaster* and *C. elegans* as well as the planarian, *Schmidtea mediterranea* (45–54). While work in planarians might suggest that mollusks can silence genes through siRNAs processed from dsRNA, further work in *C. teleta* (an annelid) shows other animals in Spiralia don’t make siRNAs (8). Within Spiralia, platyhelminths are basal to annelids and mollusks (55, 56). Consequently, if loss of siRNAs occurred before the split of annelids and mollusks, then both groups would not have endo-siRNAs. This motivates a re-evaluation of dsRNA-based gene silencing in mollusks. To establish approaches for gene silencing in these animals, we conducted a survey of the sRNAs found in mollusks. We investigated sRNA biology in 32 mollusks, with emphasis on the Eastern oyster, *Crassostrea virginica*. Results show significant similarities between annelids and mollusks. Both clearly have miRNAs and piRNAs, but there is no evidence of endogenous siRNAs. This suggests that alternatives to long dsRNA will be needed for RNAi in mollusks. Investigation of the other pathways, miRNAs and piRNAs, found that while miRNAs function similar to other animals, piRNAs appear to be restricted to specific cell types while also showing a high level of variability among individuals.

## Results

### Conservation of RNAi machinery in Mollusks

To assess available gene silencing pathways in mollusks, we characterized the small RNA biogenesis factors, Dicer and Ago/Piwi proteins, in 35 genomes (**Figure 1, Sup. Table 1).** Using annotations of genomes in NCBI and Ensembl Metazoa along with BLAST searches, putative homologs of Ago/Piwi and Dicer proteins were identified (**Sup. File 1, Sup. File 2)**. Next, using the MEGA 12 software package, multi-sequence alignments of peptides were generated to predict function through homology to well-characterized RNAi factors. Orthologs from *C. teleta, T. castaneum*, *B. mori*, *D. melanogaster, C. elegans*, *L. variegatus, S.mediterranea,* and *H. sapiens* were also included (**Figures 1A-B, Sup. Figures 1-2**). These species were chosen due to either the extensive characterization of their RNAi factors or position on the animal phylogenetic tree (57–60). Importantly, insects and flatworms (*T. castaneum*, *B. mori*, *D. melanogaster,* and *S. mediterranea)* have siAgo and siDicer. In all mollusks, miRNA and piRNA biogenesis factors were clearly identified, which was not the case for those dedicated for siRNA biogenesis (**Sup. Figure 3**).

**Figure 1.**
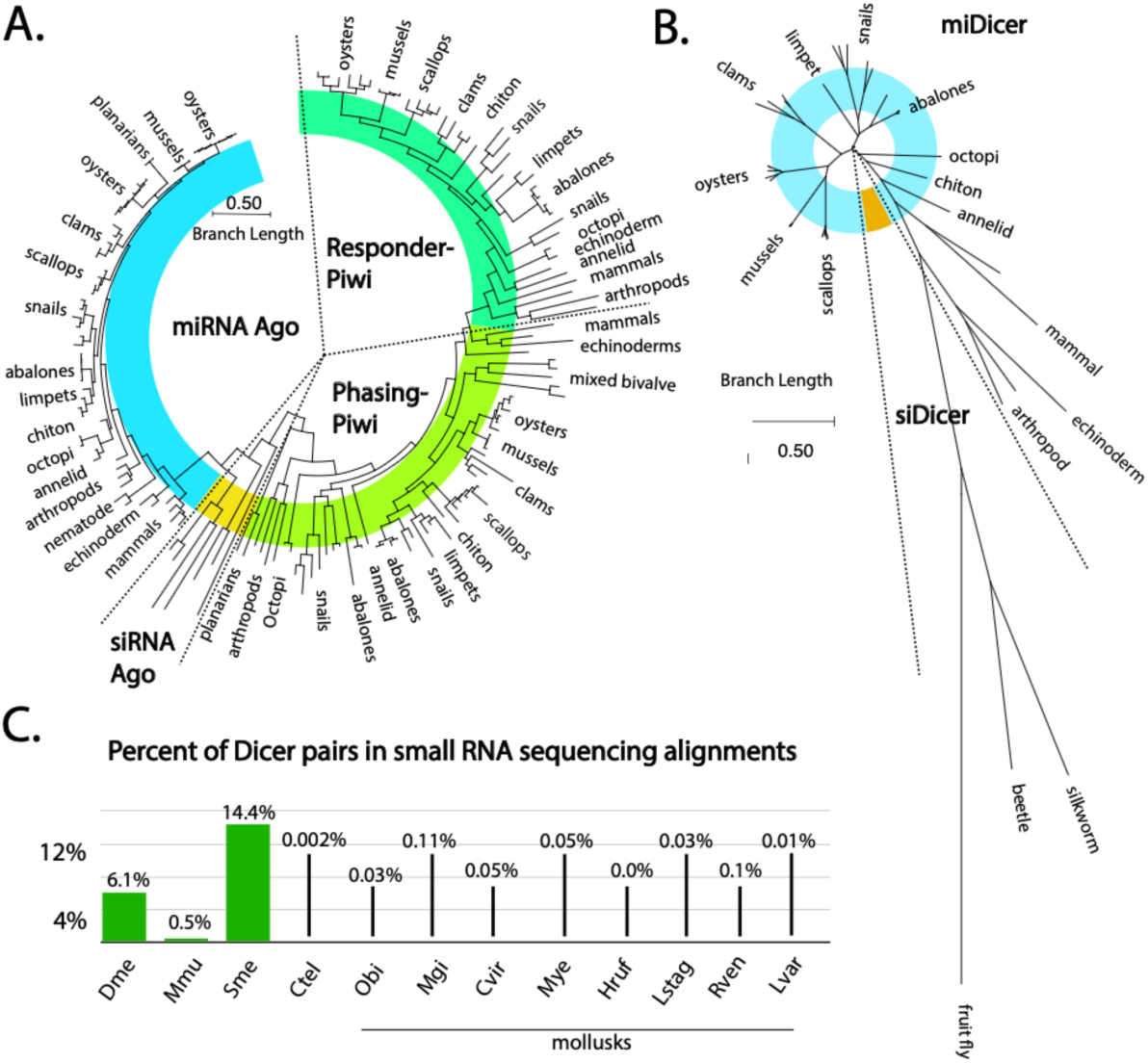
Argonaute, Dicer, and siRNA-like alignments in mollusks. **(A)** Phylogenetic relationships among the Ago/Piwi proteins of selected species. While numerous miAgo (blue), phPiwi (light green), and rePiwi (dark green) were identified in Mollusk, siAgo (yellow) was only identified in outgroup reference species: *D. melanogaster*, *B. morii*, *S. mediterranea*, *T. castaneum*, and *C. elegans*. **(B)** Phylogenetic relationship among the Dicer proteins. Numerous miRNA (blue) were identified in mollusks, siDicer was only identified in the outgroup reference species. **(C)** Percent of siRNA-like alignments found in alignments of small RNA sequencing data in *D. melanogaster*, *M. musculus*, and *S. mediterranea*, *C. teleta*, *C. virginica*, *M. gigas*, *H. rufescens*, *M. yessoensis*, *O. bimaculoides*, *L. stagnalis*, *R. venosa*, *L*. *variegatus*. The percentage indicates the number of number of small RNA reads that have overhangs consistent with RNase III processing compared to total alignments of small RNAs to the genome.

All mollusks have at least one miAgo, with several examples of group specific expansion (**Figure 1A, Sup. Figure 1**). For example, all oyster species have two miAgos that appear to be the result of duplication before their diversification. A similar situation is seen with Octopoda. There are some additional cryptic duplications seen in snails and clams. Branch lengths in the siAgo clades are very short, suggesting strong sequence conservation, as is typical for these proteins(9). Greater differences were seen in the Piwi homologs, specifically phPiwi class proteins, with duplications that occurred early in the diversification of groups. Many species have a single phPiwi; however, some bivalves, abalones, and snails have additional members that have significantly diverged sequences. Apart from the limpet *P. vulgata*, which has three phPiwis, only one additional phPiwi was observed where there was duplication. rePiwis, on the other hand, were found as single copies in all mollusk genomes. Overall, relatedness of different Ago/Piwi proteins followed phylogenetics of the clade. One exception was *G. aegis*, which likely has an accelerated rate of divergence due to extremophile adaptation. While miAgo has seen duplication, miDicer is more static, with all mollusks having only a single Dicer protein (**Figure 1B, Sup. Figure 2**). Relatedness of miDicer proteins also followed phylogenetics of the clade, with a distinct divergence between the bivalves and gastropods, and a greater divergence between Mollusk miDicers and outgroups. No mollusk siDicer-like proteins were identified that clustered with insect siDicers; instead, long branch lengths were observed, which is consistent with rapid evolution (61, 62).

To further investigate the predicted RNAi pathways in Mollusca, we performed small RNA sequencing on samples from *C. virginica* gonad, muscle, gill, and whole juvenile animals, along with muscular tissue from *H. rufescens*, tentacle tissue from *O. bimaculoides*, and gonadal tissue from the echinoderm outgroup *L. variegatus*. Public sRNA-seq data from *M. gigas*, *M. yessoensis*, *L. stagnalis*, and *R. venosa* were also included to represent groups in the phylum (**Sup. Figure 4, Sup. Table 2**). In total, ∼2.3B small reads were analyzed, with mapping rates ranging from ∼63% to ∼98%. Using alignments of these datasets, presence of an siRNA pathway was examined using an algorithm that finds pairs of reads that overlap in alignments that are ∼20-24 nt and also have 2 nt overhangs (25). Such read pairs represent potential Dicer products due their similarities to RNAs cleaved from dsRNAs. The number of these potential Dicer pairs was then divided by the total alignments from the library to calculate percent of Dicer pairs. For reference, this was also calculated for *D. melanogaster* and *S. mediterranea,* which have endogenous siRNAs, and *Mus musculus*, which has only a few endogenous siRNAs (63–65). As a negative control, we included *C. teleta*, which has no known siRNAs (**Figure 1C**) (8). In species with siRNAs, 14.4%-0.05% siRNA-like sequences were found; in sharp contrast, each Mollusk species’ results were at a minimum 5-fold lower, showing similarity to *C. teleta*. Our findings reinforce the observation that siRNAs are not made in mollusk cells. This is consistent with loss of siRNA machinery occurring before the split with annelids (8).

To understand if miRNA biology has also changed in Mollusks, we used the mirDeep2 software to annotate and document conservation patterns with the sequencing datasets (**Sup. Table 3**) (66). Many conserved miRNA families were identified across the clade, with mir-2 and mir-92 having the highest number of duplications. Numerous putative novel miRNAs were also noted in our survey (**Sup. Files 3-8**). Together, this shows that miRNA function has not significantly changed in mollusks, consistent with conservation patterns of miAgo and miDicer.

### Mollusk sRNA Population Survey

Next, to further inform gene silencing approaches in mollusks, we characterized endogenous sRNAs of 8 species for which we have sequencing data **(Figure 2, Sup. Table 2)**. First, the size distribution of all reads aligning to genomes was determined. This allows differentiation between Dicer products (miRNAs and siRNAs) that are 18-24 nt long and piRNAs, which are 26-30nt. For all mollusks, two peaks were seen that represent these two different types of small RNA **(Figure 2A-F)**. As a comparison, this was also performed on libraries from *D. melanogaster* gonads and tissue from *L. variegatus* (urchin), which show the same two peaks (**Figure 2G,H**). The relative portion of Dicer-produced small RNAs to piRNAs varies by species, suggesting divergent roles for the pathways; however, some of this variability might be attributable to libraries being created from different tissues. Some of the datasets (i.e. *C. virginica*, *L. stagnalis*) show a peak below 18 nts, which likely represent degradation products, further suggesting that inter-library differences might contribute to the distinct contours of the size distributions. Nevertheless, it is clear that in all the mollusks as well as urchins have somatic piRNAs.

**Figure 2.**
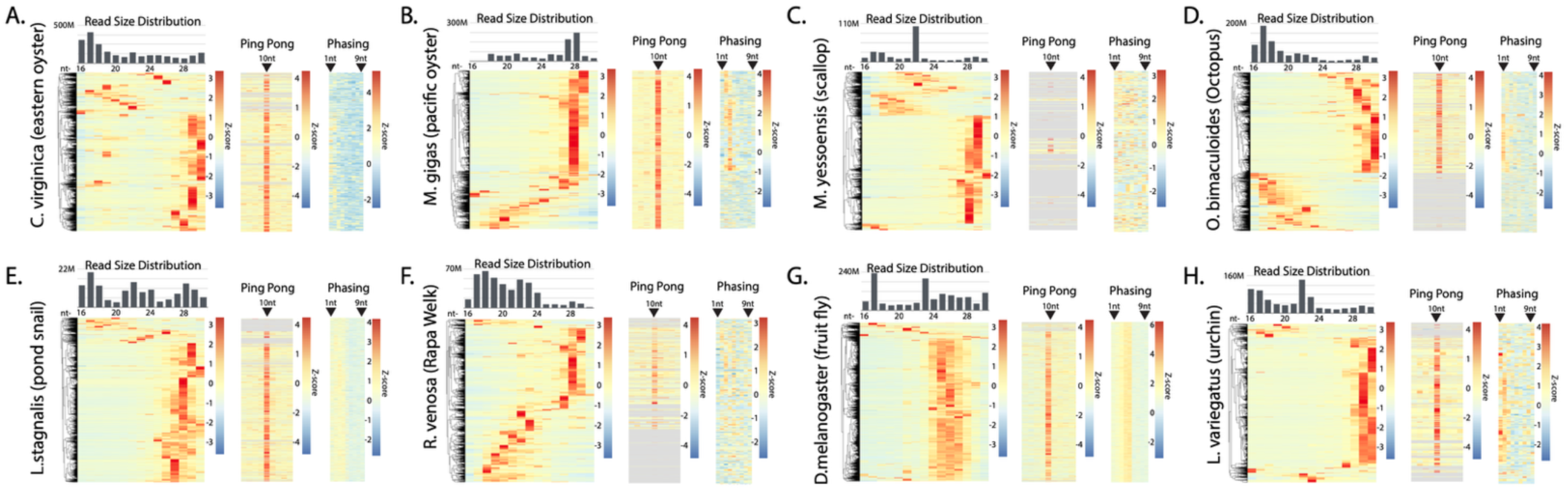
small RNA species in mollusks. Size distribution of small RNA and piRNA processing signatures in small RNA alignments found in **(A)** *C. virginica* (∼5,800 sRNA-producing loci), **(B)** *M. gigas* (∼5,400 loci), **(C)** *M. yessoensis* (∼3,800 loci), **(D)** *O. bimaculoides* (∼5,700 loci), **(E)** *L. stagnalis* (∼5,900 loci), **(F)** *R. venosa* (∼4,800 loci), (**G)** *D. melanogaster* (∼ 7,800 loci), and **(H)** the outgroup *L. variegatus* (∼3,500 loci). Each bar graph shows bulk reads across all the species’ libraries. Each heatmap (row normalized) shows the sRNA size distribution per loci. Ping Pong signature shows 10 nt overlap expression. Phasing signature is based on the distance of 1U reads to an upstream alignment. Positions 1 to 9 from the upstream alignment are shown. Order of loci in each heatmap are preserved from clustering of size distribution.

To further investigate piRNAs, we annotated small RNA-expressing loci using small read depth and merging adjacent features (8). Due to the differences in sequencing depth, appropriate thresholds were established from *C.virginica* data (**Sup. Figure 5**). By merging nearby features, accidentally splitting clusters into multiple candidate loci could be avoided. Similarly, setting a minimum read depth avoids inclusion of low confidence loci. For this analysis, *C. virginica* was used since it has chromosome-level genome assembly and is the species for which we have the greatest volume of small RNA-seq data. To determine the threshold, regions of interest were combined when within either 0, 5, 50, 500, 5000, or 50000 bp of one another. The subsequent loci of these mergers were then filtered for loci with a minimum read depth ranging from 0 to 1 million reads. We concluded that regions of interest that 1) were merged with adjacent features within 500 bp and 2) had a coverage of ≥1,000 RPM yielded both an adequate number of loci candidates (**Sup. Figure 5A**) and optimal genome coverage, which constituted 0.36% of the *C. virginica* genome (**Sup. Figure 5B**). In *C. virginica*, this methodology resulted in ∼5,800 loci-expressing sRNAs being identified (**Sup. File 7**). We then applied this coverage threshold as to identify sRNA regions in other species (**Sup. Files 10-16**).

Using the optimized annotations, we assessed the read size distribution at individual loci in the species to assess abundance of loci generating different sRNA classes (**Figure 2**). To visualize loci, values were Z-normalized and plotted as rows of a heatmap that was hierarchically sorted. For all species, including urchin and fruit fly, >50% of the loci had a bias for piRNA-sized reads. miRNA loci, in contrast appear to be a minority. Consistent with the overall size distributions, some species, such as *R. venosa*, show many loci where putative degradation products map. To further validate piRNA candidate loci, Ping Pong and phasing was assessed through calculating overlap bias and distance between reads, respectively (67). At least some Ping Pong alignments were seen in all species at loci that also showed bias for piRNA-sized reads. Similarly, phasing was detected at potential piRNA loci with the exception of *M. yessoensis* (**Figure 2H**). This is likely the result of poorly assembled genome. For example, while *C. virginica*’s genome assembly consists of only 11 contigs, *M. yessoensis*’s assembly consists of 82,659. The presence of abundant apparent piRNA expressing loci in Mollusca is similar to observations in *C. teleta*. This further reinforces the observation that mollusks and annelids share RNAi biology.

### Localization of *C. virginica* piRNAs

Most mollusks appear to have abundant piRNAs in both soma and germline, suggesting that piRNA-based gene silencing may be possible, as this has been observed with other animals with somatic piRNAs (40). To explore this possibility, we examined the expression of Piwi in tissues of *C. virginica* (**Figure 3**). Eastern oysters are often raised in aquaculture and therefore are a good candidate for biotechnology (68). We first sought to determine if similar piRNA expression was observed in both somatic and germline tissue. To accomplish this, we compared piRNA expression using samples from muscle, gill, and gonadal tissue. Selecting for only piRNA-like loci based on both their size distribution and high Ping Pong z-score, we compared loci expression in each tissue type (**Figure 3A**). Our comparison revealed distinct piRNA pathways in each of the tissue types, with some shared expression between the gill and gonadal tissue.

**Figure 3.**
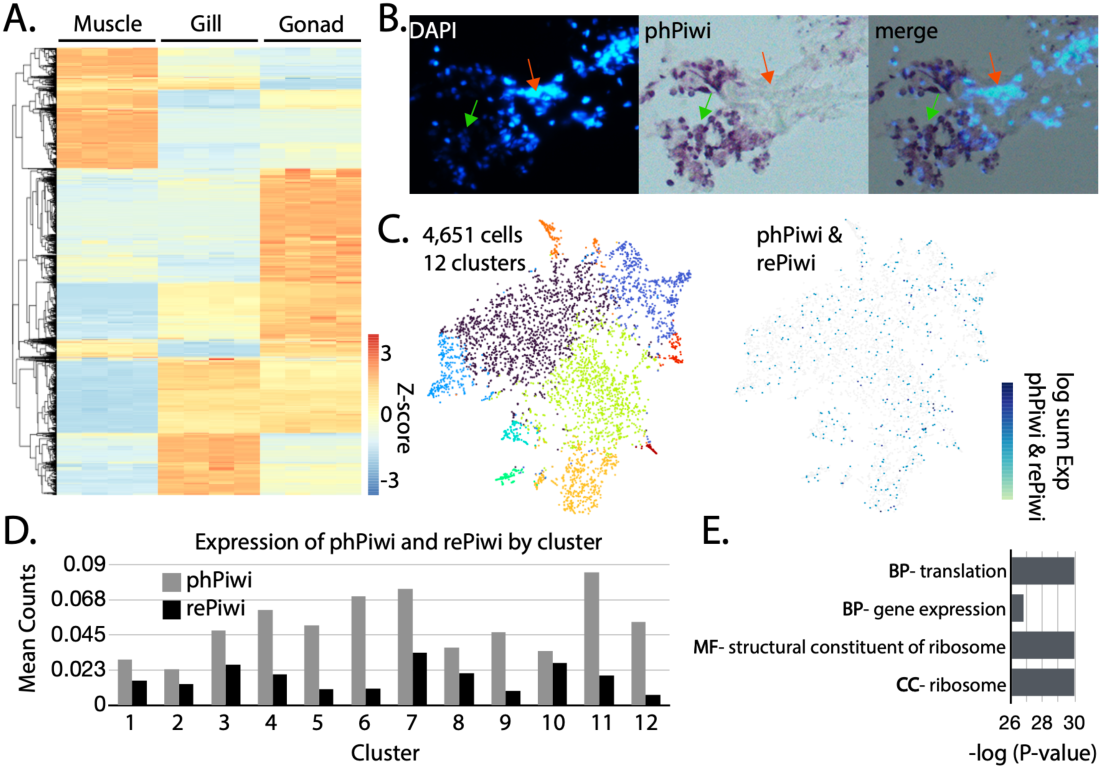
piRNA and Piwi expression in *C. virginica* tissues. **(A)** Row normalized heatmap showing piRNA (26-30 nt) expression at putative piRNA loci in *C. virginica* muscle, gill, and gonad tissue. **(B)** An overlay of DAPI staining of nuclei and colorimetric visualization of Piwi via *in situ* hybridization. **(C)** tSNA plot showing clustering of 4,651 cells into twelve clusters (left) from *C. virginica* mantle tissue. Detection of cells that express phPiwi and/or rePiwi found in the cell population. **(D)** Mean counts of phPiwi and rePiwi in the different cell clusters. (E) GO analysis of genes differentially expressed in phPiwi and rePiwi expressing cells. BP = biological process, MF = molecular function, and CC = cellular component.

While all tissues expressed piRNAs, the variability between the tissues raised the question of what cell types generate the piRNAs. To resolve if piRNAs are expressed broadly throughout tissues or if they are only present in subsets, we localized expression of *C. virginica* phPiwi using *in situ* hybridization on tissue from the mantle (**Figure 3B, Sup. Figure 6**). From this, we found that not all cells were stained for phPiwi, indicating that piRNAs are likely restricted. This observation suggested that though piRNAs are present in each tissue, they are only present in certain cell types; thus, a clearer understanding of cell identities would be needed for piRNA-based gene silencing to be realized. To characterize the identities of Piwi expressing cells, we pioneered single-nuclei RNA sequencing (snRNA-seq) in oysters using mantle tissue **(Figure 3C)**. This experiment led to recovery of 4,651 cells which clustered into 12 groups. Oyster mantle has been described on the histological level, but not at all on the cellular (69). This makes it challenging to understand the identities of cells; however, when searching for *C. virginica* phPiwi and rePiwi transcripts within the clusters found, they did not map specifically to any group.

Looking at counts of the phPiwi and rePiwi within the clusters, we found some variability, with phPiwi having a higher expression level, though transcripts were clearly found in all **(Figure 3D)**. Together, this shows that Piwi expressing cells are not a well-defined cell type in a tissue, but rather a subset of many cell identities.

To understand the unique features of Piwi expressing cells, differential expression was performed for these cells relative to the overall population. Roughly 90 genes were found to be differentially expressed, nearly all of which were downregulated in the Piwi cells. GO analysis was then performed on the gene set, which showed highly significant enrichment for genes involved in translation **(Figure 3E)**. Identities of the genes included both initiation factors and components of the ribosome itself. All these genes related to translation in the gene set were downregulated, suggesting that Piwi expression co-occurs with quiescence. Such a state is observed in stem cells, which may be involved in the maintenance of oyster tissue (70). If the piwi positive cells are in fact stem cells within each cell type, this would align with observations in *C. teleta* (71). These stem cells may be involved in regeneration and maintenance of the tissue. Alternatively, the cells could be experiencing a state of stress and Piwi expression is a component of that response (72).

### Role of piRNAs in mollusk regeneration

Work in *C. teleta* demonstrated that Piwi expression is present in regenerating tissues and other proliferative cells, and the work here in oysters suggests a similar biology (71). To investigate if piRNAs have a consistent function that relates to regeneration, we investigated piRNA expression in intact and regenerating tissues in *C. teleta, C. virginica, L. stagnalis, O. bimaculoides, and L. variegatus* (**Figure 4A,B**). For each species, somatic tissues were harvested (**Sup. Table 2**). All data, excluding those from *L. stagnalis* (Bioproject PRJNA664475), were generated as part of this study. Consistent with observations in *C. teleta*, we found significant changes in piRNAs annotated in prior work expressed in intact vs regenerating tissues in this annelid (**Figure 4A**). Comparison of individual samples revealed high similarity between the tissue types (**Figure 4B**). Unexpectedly, an opposite outcome was seen in the oyster piRNA loci annotations shown in Figure 3. Only a single piRNA locus showed differential expression and it was upregulated by ∼1.4 log2 fold change. The difference extended to clustering via PCA, where samples from the same individual showed more similarity than tissue types (**Figure 4B**). This situation was also seen in *L. stagnalis*, with only a handful of differentially expressed piRNA loci. Unfortunately, the public dataset metadata did not indicate whether intact and regenerating samples were from the same individual; however, based on sample clustering, some may have been acquired from the same animal. *O. bimaculoides* data was more similar to *C. teleta*, with many significantly differentially expressed piRNAs. Unlike other species examined here, samples were collected from a single individual at different sites (tentacle tips). As part of this study, we also examined piRNA expression during regeneration in *L. variegatus* as an outgroup.

**Figure 4.**
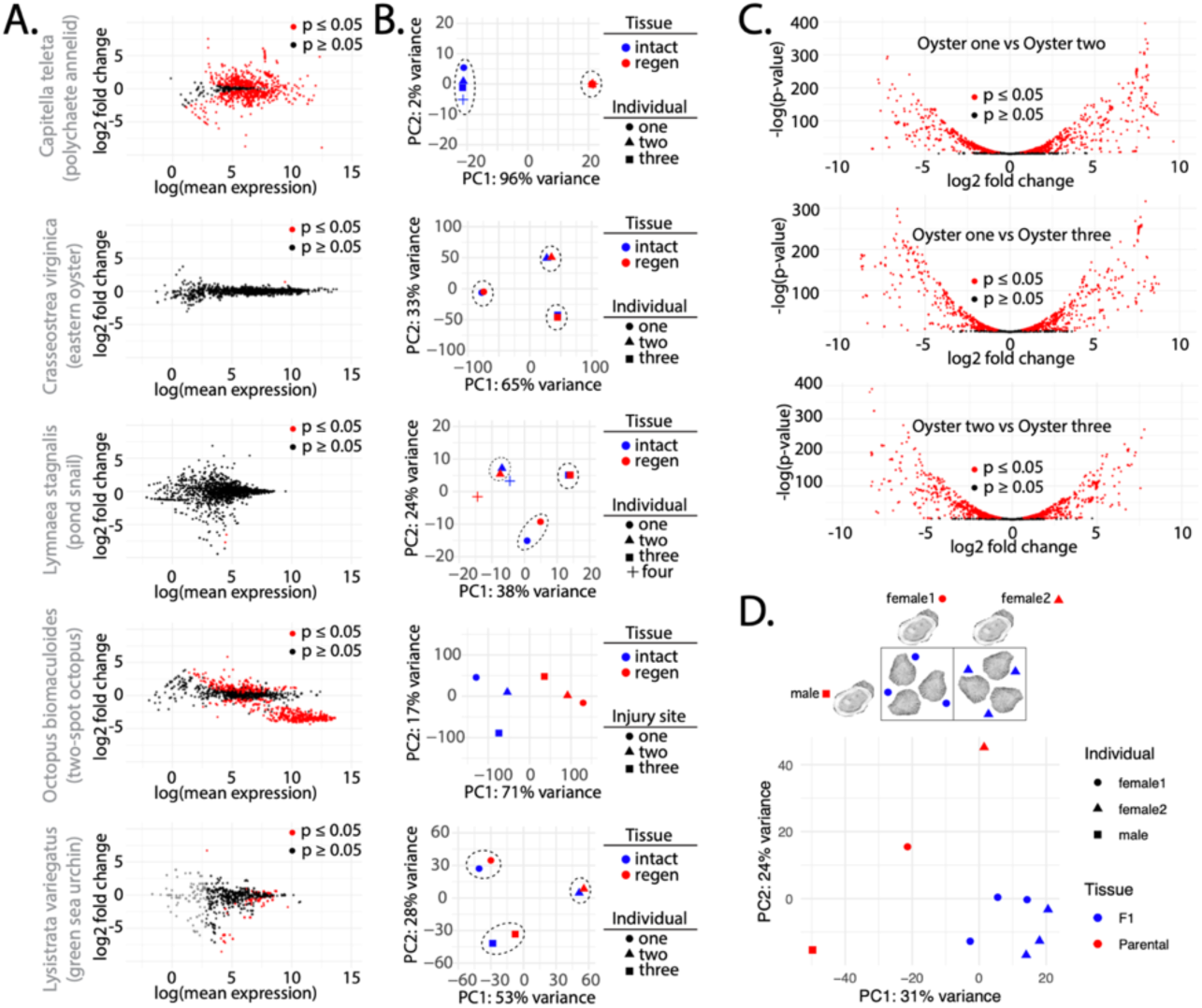
piRNA expression during regeneration and comparison of expression between parental and F1 animals. **(A)** Scatterplots showing piRNA differential expression in uninjured tissue and regenerating tissue in *C. teleta*, *C. virginica*, *L. stagnalis*, *O. bimaculoides*, and *L. variegatus*. Black dots are not significant (p value ≥ 0.05), while red dots are significant (p value ≤ 0.05) **(B)** PCA of piRNA expression in each species’ individual intact (blue) and regenerating tissues (red). Dashed lines highlight closely clustering individuals. **(C)** Volcano plots showing the differential expression of piRNAs between datasets from individual animals. Black dots are not significant (p value ≥ 0.05), while red dots are significant (p value ≤ 0.05). **(D)** PCA of piRNA expression in parents and F1 easter oysters. Breeding scheme shown above where sperm from a single male (square) was used to fertilize eggs from two different females (circle and triangle) to yield two sets of juvenile oysters. Relation of juvenile oysters to which mother shown by matching shape of the data point (circle and triangle).

Interestingly, the urchin data showed similarity to *C. virginica* and *L. stagnalis*. This suggests that the highly individualized piRNA expression that we see in oysters and snails might be the ancestral state, and that the situation in *C. teleta* is derived. The octopus data suggest that piRNA expression may likely change during regeneration in mollusks; however, individual differences in piRNA abundance might exceed the changes that occur in response to tissue damage.

When comparing the differences in expression between the individual oysters–comparing intact and regenerating samples from one oyster to another– we found highly divergent patterns of piRNA expression (**Figure 4C**). These animals were all raised in a single hatchery from the same brood stock. Despite this, the number of loci showing differential expression exceeded 55% of the total number of piRNA annotations. To investigate the origin of the disparate piRNA expression seen between animals, sperm from one male was used to fertilize eggs from two females, collecting both parental and F1 tissues (**Figure 4D**). Here, one male oyster was used with two females. Offspring were then allowed to mature for three months to a juvenile stage. The breeding scheme was designed so that if F1s showed similarity to a parent, this would indicate the basis of an individual’s piRNA configuration. When comparing variance in piRNA expression via PCA, we found the juveniles cluster together, while the parents were quite distant from their offspring and each other. On the PC2 axis, offspring showed closeness to the father, while both mothers were further from their respective children. This is a little surprising given that piRNAs have been found to be inherited maternally in an extra-nuclear fashion (73). These results suggest that piRNA repertoire in oysters is contributed by the genetics of both parents, and that as animals age, expression may become increasingly divergent.

## Discussion

Exploration of Mollusk small RNA pathways found many miRNAs and piRNAs but no evidence of siRNA pathways. Mollusk piRNAs have a distinct biology relative to ecdysozoans and vertebrates with common somatic piRNAs that appear to be restricted to only a portion of cells within a tissue. piRNA expression patterns are also highly individualized and are partially defined by inheritance. Moreover, unlike other animals it appears that piRNA expression is determined equally by maternal and paternal inheritance. Together these observations prescribe the application of RNAi-based technology in mollusks.

Mollusk fisheries in the United States face significant challenges, highlighted by a 93% decline in eastern oyster landings due to the collapse of wild harvesting (74–80). To combat this decline, aquaculture has become the predominant method of propagating many species (68). This provides an opportunity for biotechnology to induce desirable traits such as robustness to environmental stress or disease resistance. This does raise some ethical and environmental concerns with modified animals being placed in the environment to fully mature. Many oyster hatcheries produce triploid offspring, which are considered sterile and therefore have a reduced concern for spread of modified alleles or transgenes into wild populations (81, 82). Many challenges exist for these approaches, including isolation of genetically modified brood stock from wild populations or delivery of large molecules such as Cas9-gRNA complexes in aquaculture facilities. These limitations could be addressed with RNAi technology as it does not lead to permanent genomic changes and can be applied with large scale RNA synthesis to oysters during early developmental stages while they are being raised in large volumes of seawater (81–83).

Gene silencing in invertebrates is typically triggered by delivery of dsRNA to exploit the siRNA pathway (84, 85). Here we show that mollusks, and likely the entire Lophotrochozoan clade, does not have this mechanism. Thus, dsRNA-mediated RNAi is not a viable means of eliciting RNAi in Mollusca. In contrast, many piRNAs are present in Mollusca. Though production of ectopic piRNAs in somatic tissues have been demonstrated in insects for gene silencing such an approach would also not be appropriate in mollusks. In this study we observe that Piwi proteins are not present in all cells within tissues. (**Figure 3B-C**) (40). Gene expression profiles in Piwi-expressing cells suggest that these are likely stem cells which piRNAs having a role in maintaining, possibly through ensuring genome integrity by suppressing of transposable elements (38, 86). Similar stem cell restriction of Piwi and piRNAs has been observed in *C. teleta* and planarians (71, 87, 88). Thus, mollusks seem to share a role for piRNAs with members of the larger spiralian clade. This lack of ubiquitous piRNAs in tissues raises concerns about the usefulness of piRNA-mediated RNAi in *C. virginica* and likely the broader Mollusca phylum. Instead, the most viable means of inducing RNAi in mollusks will likely be through exploiting the miRNA pathway, as is the basis of gene silencing in humans (89–91).

In addition to informing technology development, the results of this study also provide insights into the fundamental biology of gene expression in mollusks. Studies have found that on the population level oyster genomes are highly polymorphic (92). This is likely a consequence of the need for sessile animals, such as oysters, to be phenotypically plastic and respond to environmental conditions. Indeed, shell formation is a complex process that generates varying external morphologies (e.g. size, shape) (93). The increased genetic variability in these animals also seems to lead to proliferation of deleterious alleles. Investigation of the Pacific oyster genome found evidence for non-Mendelian inheritance that likely compensates for a dominant high genetic load (93). Additionally, heterosis has been observed in oysters, suggesting significant individual genetic differences (94). Other studies have found significant chromosomal abnormalities accumulating in oyster tissues, suggesting the negative alleles are also accompanied by genomic instability (95). This situation informs the likely role and abundance of piRNAs in oysters. By expressing these RNAs in a stem cell-like compartment, better genome integrity in this cell type could help preserve cell viability during growth or regeneration. Bivalve genomes seem to be routinely remodeled by transposons (96). The expansion of parasitic genetic parasites could be the basis of genetic variability seen in oysters and reinforces the importance of piRNAs in balancing transposon activity throughout the animal’s tissues.

## Methods

### Animal Husbandry

*C. virginica* specimens were obtained from the Thad Cochran Marine Aquaculture Center in Ocean Springs, Mississippi. Specimens were kept in a 200-gallon tank with biofilters and a UV sterilizer, and we maintained ∼20 ppt salinity and regularly monitored levels of pH, ammonia, nitrate, and nitrite. For feeding, we moved the oysters into a separate container with sea water and fed them ∼30 mL (∼6 x 10^10^ algae cells) of Shellfish Diet © 1800 every other day for ∼4 hours each feeding.

*C. teleta* worms were obtained from the lab of Dr. Elaine Seaver at the Whitney Laboratory for Marine Bioscience at the University of Florida. Housed in a 20° C growth chamber, our specimens were kept inside containers with 100 ml of estuary mud and 200 ml of salt water. Each container held ∼20 worms, evenly divided between males and females. The boxes were monitored for the presence of swimming larva, which were then transported to new containers. The worms were fed weekly by additional estuarine mud.

*L. variegatus* specimens were obtained from Gulf Specimen Marine Lab in Panacea, Florida. Animals were kept in a 50-gallon tank with biofilters and large rocks to simulate the conditions of a reef, with ∼25 ppt salinity. The urchins were fed every other day one-third of dried nori algae sheets bought from a local market.

A female *O. bimaculoides* was obtained from Marine Biological Laboratory. The octopus was housed in a 70-gallon tank with biofilters and ∼36 ppt salinity. A chiller was attached to the pump to keep the water at ∼18° C. Clay pots, rocks of varying sizes, and plastic blocks were placed inside the tank for shelter and entertainment. The octopus was fed thawed shrimp daily. A live specimen of *H. rufescens* (red abalone) was purchased from Monterey Abalone Company and sacrificed immediately to acquire samples.

### Animal Injury and Regeneration

#### Crassostrea virginica

Three adult oysters of similar size were selected and injured by first shaving their shell to create a smooth surface followed by drilling a hole through the shell into the mantle region. The oysters were allowed two weeks to partially regenerate their shells, which we confirmed by observing a thin layer of secreted shell covering the drilled holes. After two weeks, we extracted proximal mantle tissue directly underneath the injury site to serve as the regenerative sample. We also collected distal tissue from the opposite side of the mantle to serve as the control. This protocol was based on a similar experiment that observed miRNA and gene expression in repair and biomineralization in freshwater snails (97).

#### Capitella teleta

Worms were placed under a dissecting microscope and amputated at the 12^th^ body segment using forceps. They were then placed in 20 mL of filtered salt water and allowed to recover for 24-48 hrs. After the recovery period, they were placed in containers with mud and salt water. The worms were checked every other day for growth and survivability, and the regenerated tissue was collected after two-three weeks.

#### Lytechnius variegatus

For urchin amputations, each individual was injured using a pair of fine scissor and nail clippers, about half of the urchin’s spines were cut very close to the test (hard shell). Once the tube feet were exposed, a pair of fine scissors was used to cut the tube feet. The urchins were placed back into the tank and allowed to regenerate for one week. After they had sufficient time to regrow some of the tube feet, they were placed back into smaller containers, and the process was repeated. The previously injured half of the urchin was collected from the container using a pipette and strained using a 40 µm cell strained to remove salt water and retain the tube feet. The unharmed half of the urchin went through the same process and was used as control.

#### Octopus bimaculoides

The octopus used in this study was handled following ethical considerations established by the European Parliament and the Council of the European Union. Before injury, it was anaesthetized using ethanol. 0.25% increments v/v of ethanol were added every two minutes to a holding container until a final concentration of 1.5% EtOH was reached. After the animal was under complete anesthesia for ∼14 minutes, the tips (∼2 cm) of three tentacles were amputated using a sterilized razor blade and collected as control samples. After allowing the tentacles to regenerate for three days, the above protocol was repeated to collect the blastemas that had formed. The animal was euthanized using the same approach as anesthesia until the EtOH concentration had reached a lethal concentration.

### Breeding and Offspring Collection

A male and two female oysters were exposed to heated seawater (32^°^ C) to induce gamete release. These parental animals were sacrificed and their muscle, mantle, gill, and gonad tissues collected. Eggs from each female were fertilized with sperm from the male, with the two crosses cultured separately. The resulting embryos from each cross were cultured to the pediveliger stage, harvested, and set on microcultch to create single set oysters. Post-set (seed) were cultured in upwelling nursery systems for ∼3 months. Several of the offspring from each of the two crosses were sacrificed and their entire bodies used for RNA extraction.

### RNA Extraction and Sequencing

All harvested tissues were placed in 700 µL of TRizol LS and 300 µL of nuclease free water. Addition of water was necessary to balance excess salts present in the tissues of these marine animals to make samples compatible with the TRizol method. After tissue homogenization, samples were processed following manufacturer protocols, and the resulting isolated RNA resuspended in 30 µL of nuclease free water. Total RNA samples were sent to the University of Mississippi Medical Center Genome Core, which prepared libraries using the Illumina small RNA TruSeq library construction kit and using an Illumina NextSeq2000.

### Phylogenetic analysis

We searched NCBI for Dicer and Ago/Piwi annotations in Mollusca species. After identifying numerous annotations, we further searched for homologs using Ensembl Metazoa and NCBI BLAST searches. We then imported our resulting Dicer and Ago/Piwi protein sequences into Molecular Evolutionary Genetic Analysis Version 12 (MEGA12), which utilizes Multiple Sequence Comparison by Log-Expectation (MUSCLE) for phylogenetic analysis.

### Small RNA Analysis Pipeline

The following genome sequences and annotation files were acquired from NCBI’s GenBank database: *C. virginica* (GCA_002022765.4), *M. gigas* (GCA_963853765.1), *Haliotis rufescens* (GCA_023055435.1), *Lytechinus variegatus* (GCA_018143015.1), *Octopus bimaculoides* (GCA_001194135.2), *Rapana venosa* (GCA_028751875.1), and *Lymnaea stagnalis* (GCA_900036025.1). Genome sequence and genome annotation files for *C. teleta* were acquired from Ensembl Metazoa (Capitella_teleta_v1.0). We clipped the adaptors off the resulting high-throughput sequencing data using CutAdapt and then used bowtie and samtools to map and process alignments to respective genomes. Using a python-based algorithm, we screened for 15-30nt sRNAs that had 2 nt overlaps (indicating Dicer signature found in siRNAs) or 10 nt overlaps (indicating Ping Pong signature found in piRNAs) (98). piRNA loci were analyzed for phasing using an approach described by the piPipes (67). We also generated a read counts table that could be visualized using ggplot2 and pheatmap. DESeq2 was used to compare the piRNA expression of individual intact tissues to one another.

### *In Situ* Hybridization and Single-Nuclei Sequencing

To localize *C. virginica* PIWIs, we harvested mantle, gill, gonad, and muscle tissue. After freezing the extracted tissues in OCT compound, we used a cryostat to section and mount 5nm tissue on Plus Gold slides. To generate a phasPiwi probe, we first cloned a portion of phasPiwi via PCR using phasPiwi-specific primers (Forward: ATGTCAGGTAGAGGAAGAGCTCGTG; Reverse: TCAAGCTATGCATCCAACGCGTTGGGTTTGTTCACATGTCCAAGCTTGC). The ThermoFisher Phire Hot Start II kit was used, and the samples cycled 35 times, with denaturation for 5s at 98°C, annealing for 15s at 60°C, and denaturation for 60s at 72°C. Following PCR amplification, T7 in vitro transcription was used to create a probe labeled with Digoxygenin. We then performed *in situ* hybridization as previously described (99). For single nuclei sequencing, mantle tissue freshly embedded in OCT was sent to the genomics core at the UMMC for single-nuclei sequencing using a Chromium X with the Chromium Next GEM Single Cell 3’ Kit v3.1 followed by sequencing on an Illumina NextSeq 2000.

## Acknowledgments

This work was supported by NSF 1845978. By the Mississippi INBRE (P20GM103476). Computational resources were provided by NSF: ACI 1626217. The work performed through the UMMC Molecular and Genomics Facility is supported through the Molecular Center of Health and Disease (P20GM144041) and the Obesity, Cardiorenal and Metabolic Diseases-COBRE (P30GM149404).

---

**Supplementary Figure 1:**
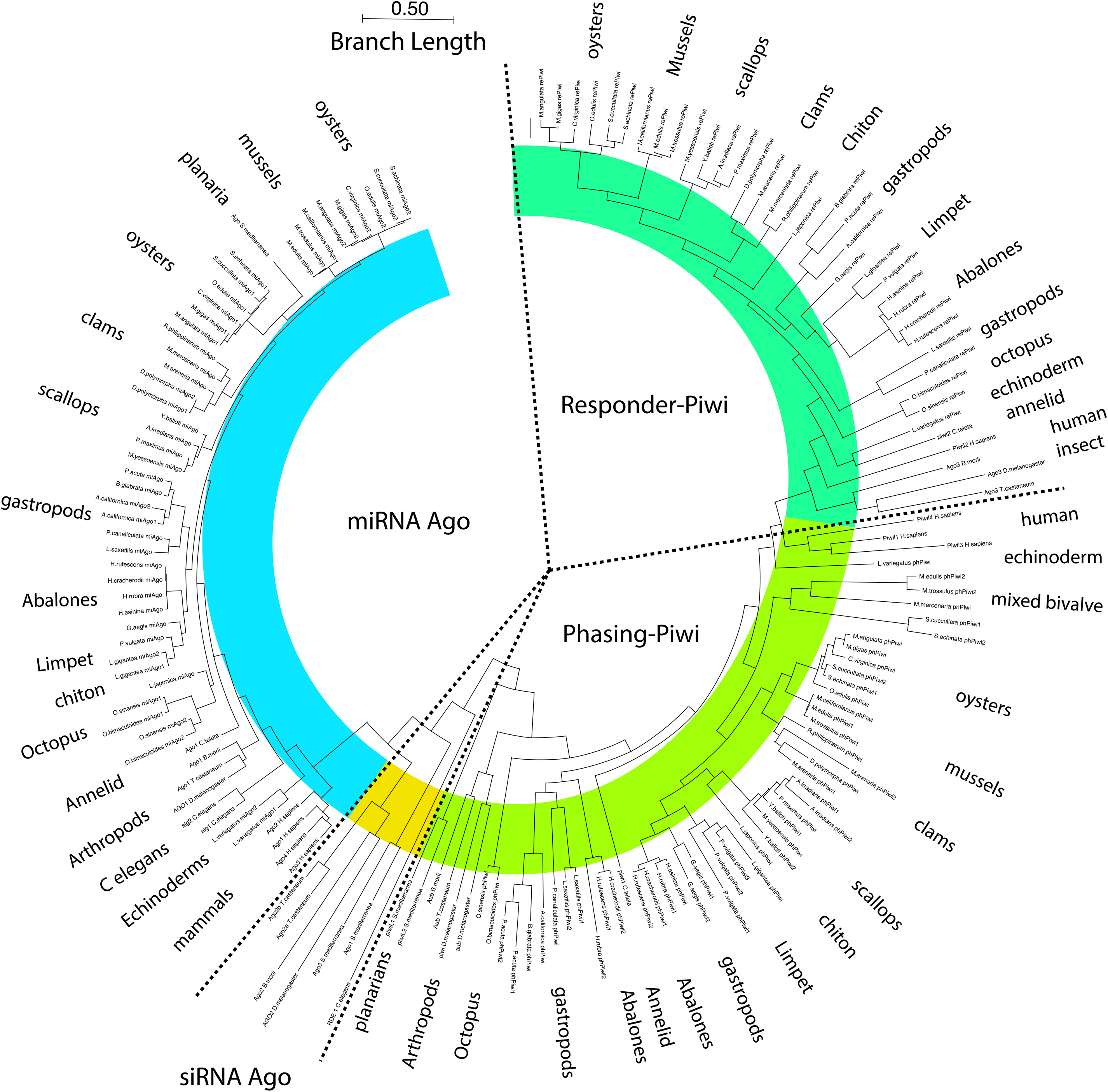
Phylogeny Analysis of AGO/PIWI Proteins in Mollusca. Comprehensive tree showing phylogenetic relationship among the AGO/PIWI proteins of different species. While numerous miAGO (blue), phPiwi (lime green), and rePiwi (pastel green) were identified in Mollusca, siAGO (yellow) was only identified in outgroup reference species: *D. melanogaster, B. morii, S. mediterranea, T. castaneum, and C. elegans.*

**Supplementary Figure 2:**
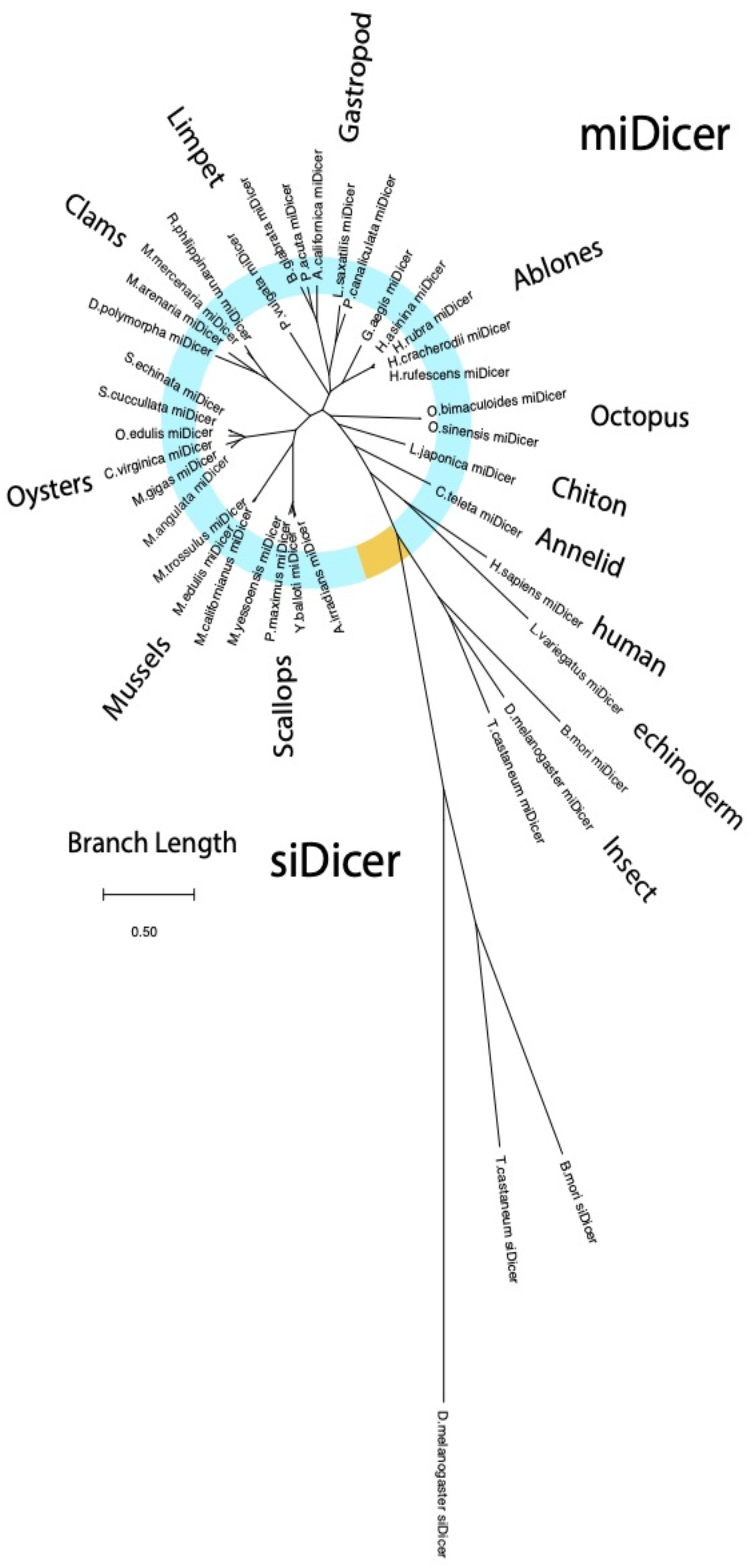
Phylogeny Analysis of Dicer Proteins in Mollusca. Phylogenetic relationship among the Dicer proteins of different species. The Dicer1 reference species (blue) are *L. stagnalis*, *C. teleta*, *H. sapiens*, *D. melanogaster*, and *C. elegans*. The siDicer reference species (yellow) are *D. melanogaster*, *Tribolium castaneum*, and *Bombyx mori*. Phylogenetic analysis demonstrates a correlation between all known Mollusca Dicer and known miDicer proteins rather than a correlation with siDicer proteins.

**Supplementary Figure 3:**
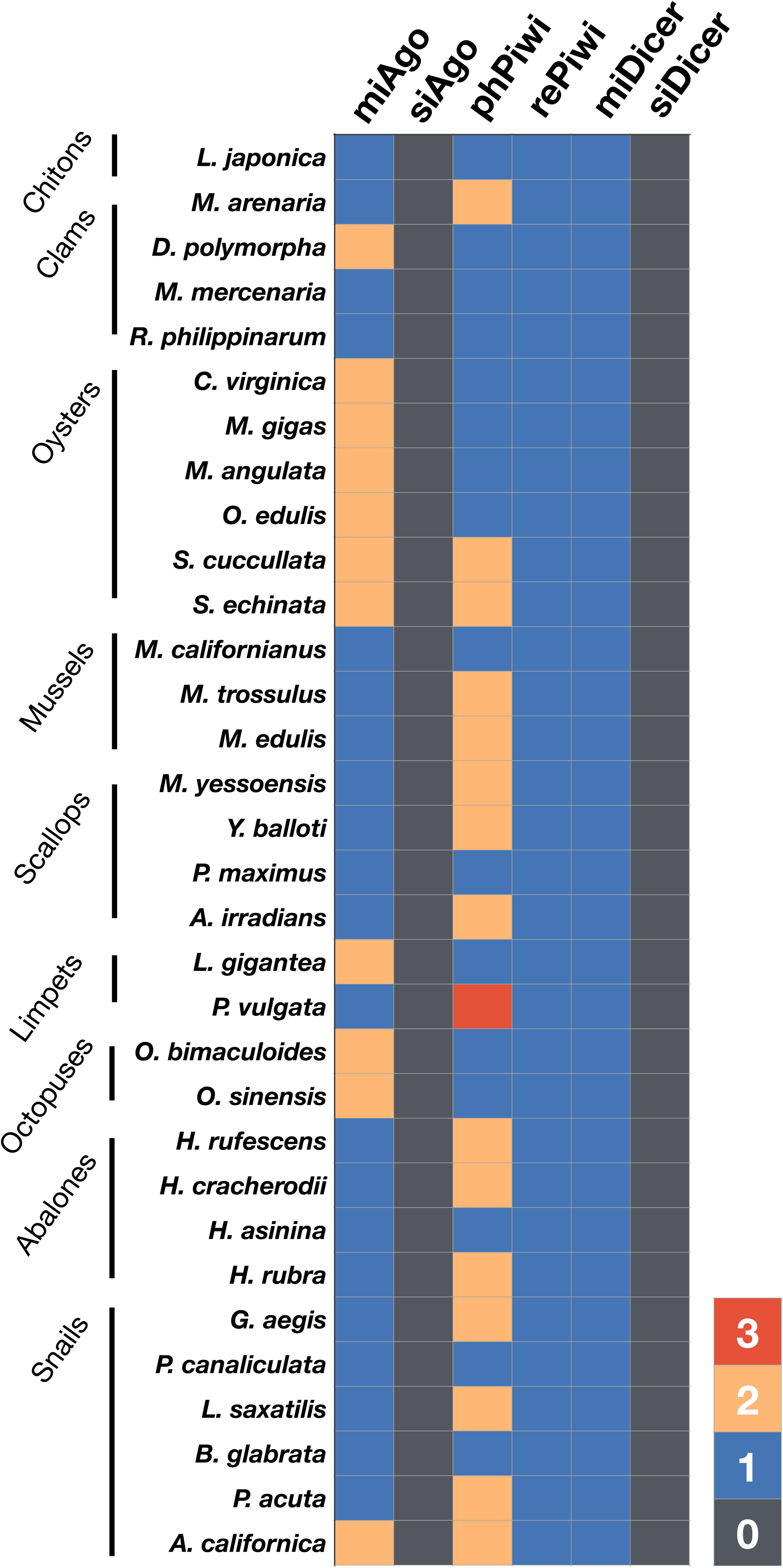
sRNA proteins, siRNAs, and miRNAs in Mollusca. An overview of sRNA-associated proteins in mollusks, sorted phylogenetically. While numerous Ago and Piwi proteins were identified, neither siAgo nor siDicer proteins were observed.

**Supplementary Figure 4:**
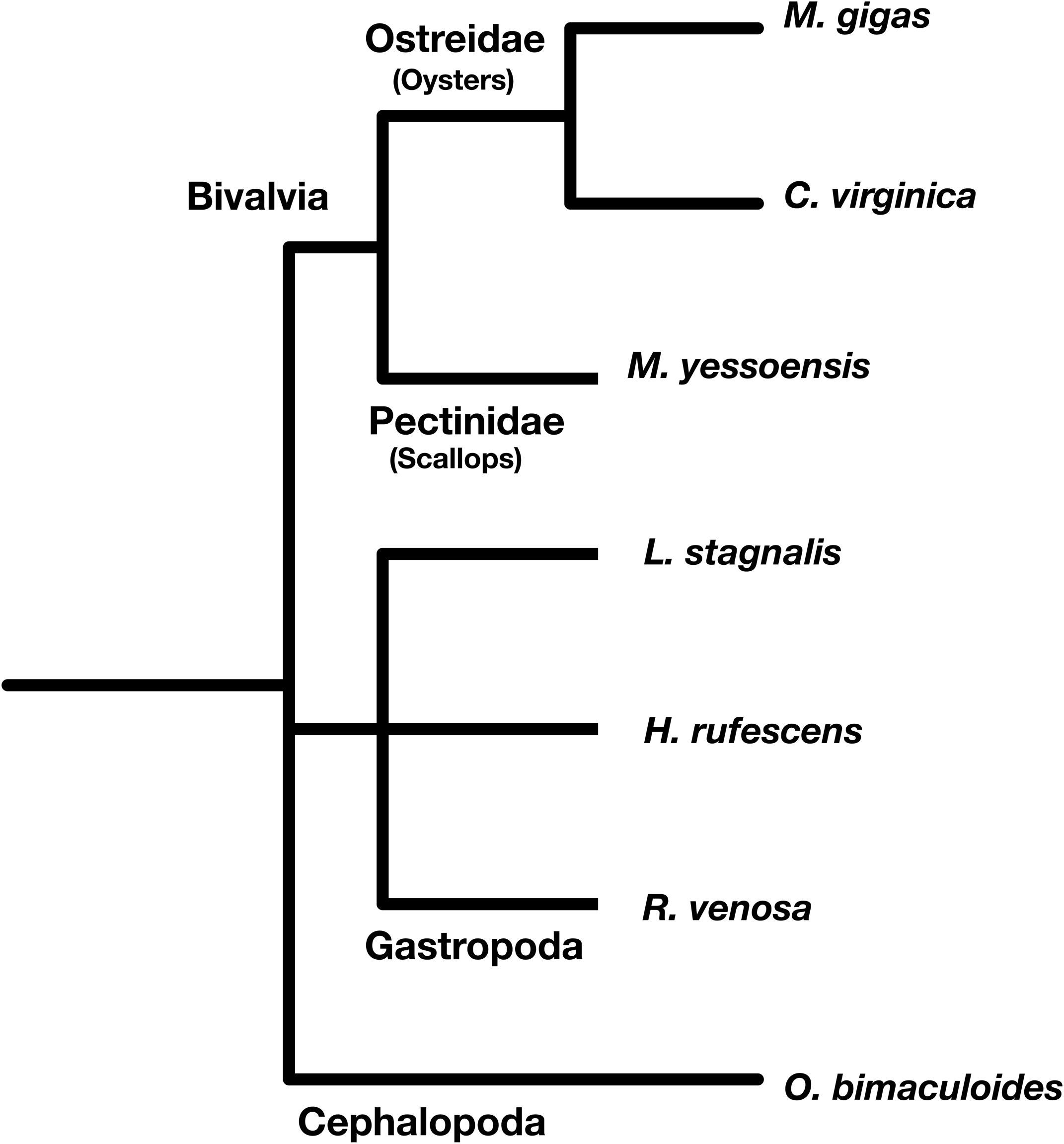
Phylogeny of Mollusca Analyzed. Phylogenetic tree within the Mollusca phylum indicating the species whose sRNA-seq data we examined in this project: *M. gigas*, *C. virginica*, *M. yessoensis*, *L. stagnalis*, *H. rufescens, R. venosa*, and *O. bimaculoides*.

**Supplementary Figure 5.**
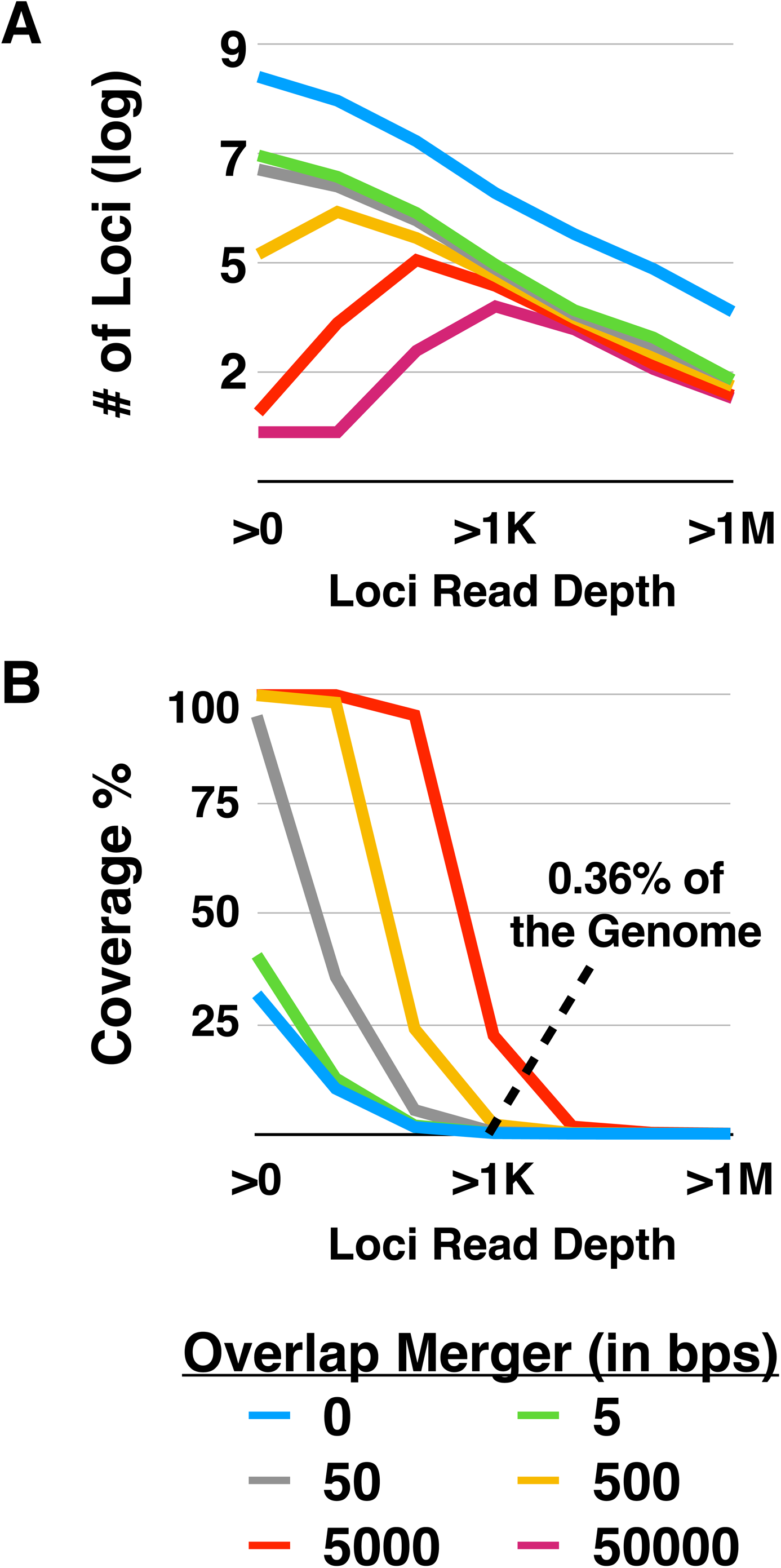
sRNA Thresholding sRNA of *Crassostrea virginica*. An optimal genome coverage threshold was determined by plotting loci read depth compared to the merging of read overlaps. **(A)** The number of loci (in log) identified when sequential overlap mergers (lines) were plotted against an increasing minimum of read depths per loci (x axis). **(B)** The loci’s total genomic coverage when sequential overlap mergers (lines) were plotted against an increasing minimum of read depths per loci (x axis). Loci that were merged with other regions of interest within 500 bp (yellow line) and had a minimum read depth of 1,000 appear to represent the median in both the number of loci present and overall genomic coverage.

**Supplementary Figure 6:**
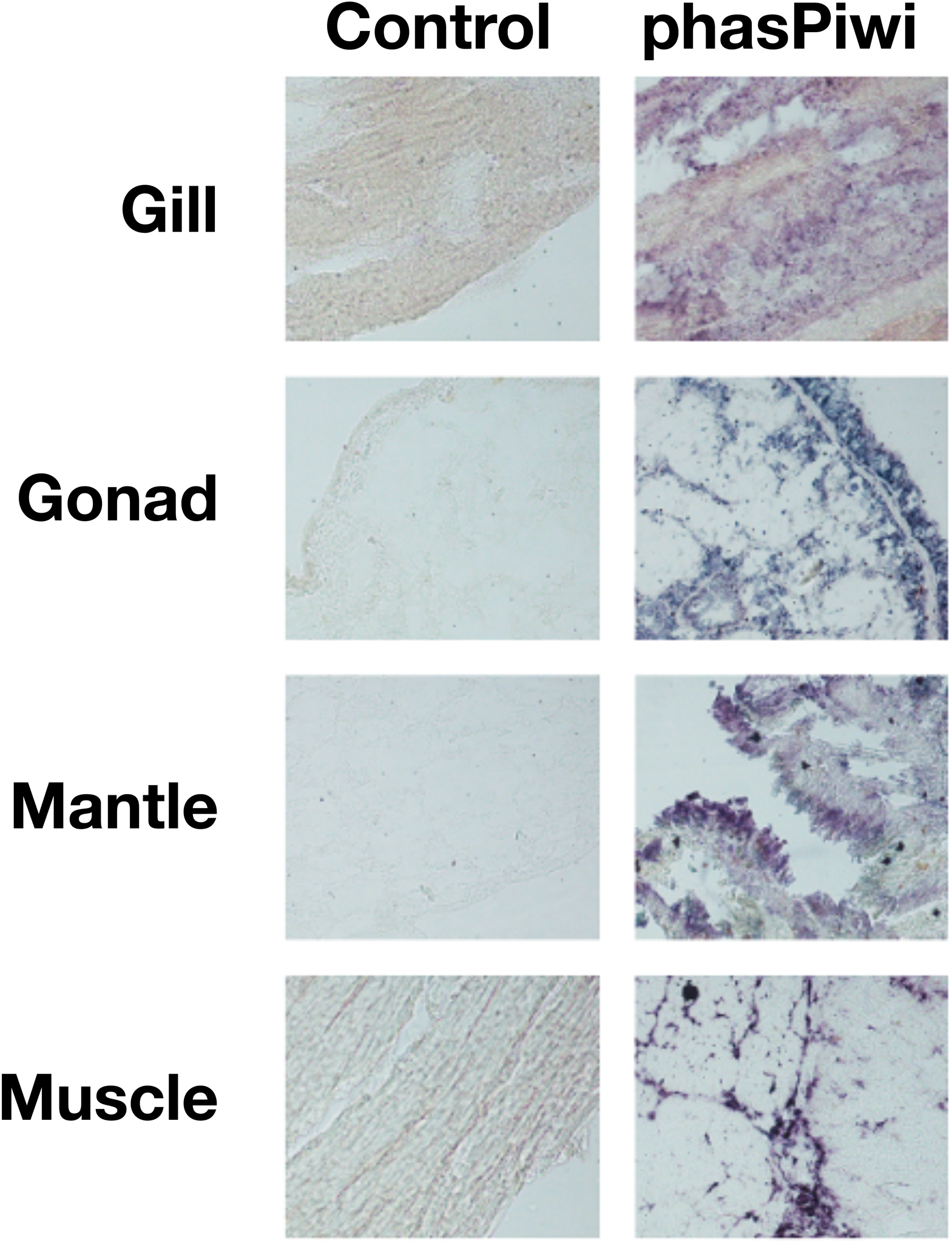
*In Situ* Hybridization of *C. virginica*. Visualization of phas-piRNAs in various *C. virginica* tissue types. The purple represents piRNAs found in cells within each tissue.

**Supplemental Table 1.**
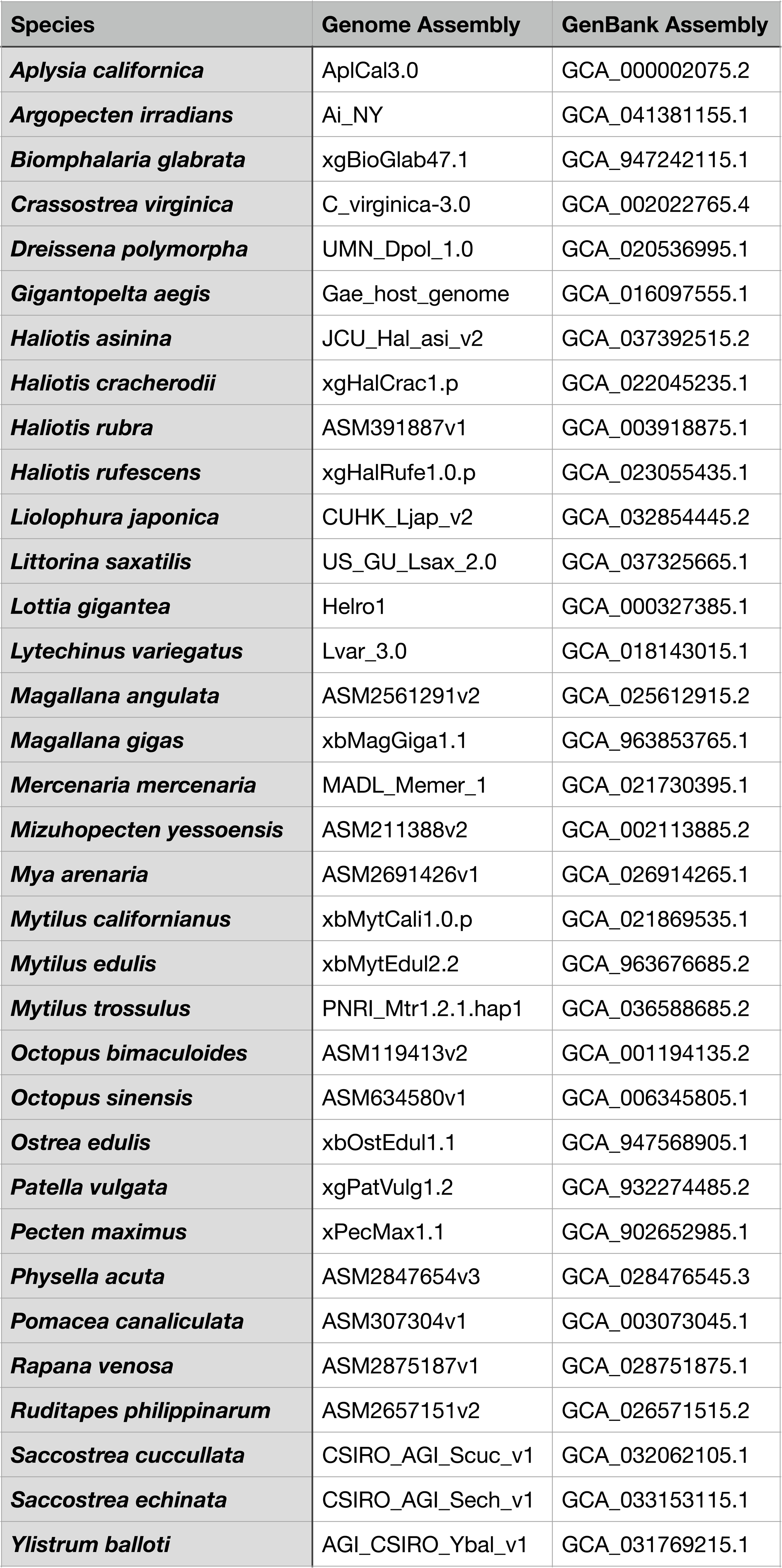
Overview of Genomes. Summary of NCBI Mollusca (and *L. variegatus*) genomes used in data computation.

**Supplemental Table 2.** Overview of Data Libraries. Summary of libraries used in data analysis of both Mollusca and outgroup species.

| Species | Tissue | SRA | Reads | Species | Tissue | SRA | Reads |
| --- | --- | --- | --- | --- | --- | --- | --- |
| *C. virginica* | INH_Male_Gonad | SRR33418593 | 51703409 | *L. variegatus* | Gonad_Ctrl1 | SRR33418561 | 40639852 |
| *C. virginica* | INH_Female1_Gonad | SRR33418592 | 59187962 | *L. variegatus* | Gonad_Ctrl2 | SRR33418560 | 31498024 |
| *C. virginica* | INH_Female2_Gonad | SRR33418581 | 50650416 | *L. variegatus* | Gonad_Ctrl3 | SRR33418558 | 48932488 |
| *C. virginica* | INH_Child1B | SRR33418570 | 49392863 | *L. variegatus* | Gonad_Reg1 | SRR33418557 | 51016857 |
| *C. virginica* | INH_Child1C | SRR33418559 | 52729896 | *L. variegatus* | Gonad_Reg2 | SRR33418556 | 44795701 |
| *C. virginica* | INH_Child1E | SRR33418549 | 49693994 | *L. variegatus* | Gonad_Reg3 | SRR33418555 | 45765823 |
| *C. virginica* | INH_Child2F | SRR33418548 | 42504756 | *L. variegatus* | INH_Male_Gonad | SRR33418554 | 136816008 |
| *C. virginica* | INH_Child2H | SRR33418547 | 52188423 | *L. variegatus* | INH_Female1_Gonad | SRR33418553 | 131933301 |
| *C. virginica* | INH_Child2I | SRR33418546 | 49327186 | *L. variegatus* | INH_Female2_Gonad | SRR33418552 | 115282154 |
| *C. virginica* | AdductorMuscle1 | SRR33418545 | 9448493 | *L. variegatus* | INH_ChildM1F1 | SRR33418551 | 134202757 |
| *C. virginica* | AdductorMuscle2 | SRR33418591 | 9348118 | *L. variegatus* | INH_ChildM1F2 | SRR33418550 | 187061029 |
| *C. virginica* | AdductorMuscle3 | SRR33418590 | 9665186 | *O. bimaculoides* | Tentacle_Ctrl1 | SRR33418569 | 56141057 |
| *C. virginica* | AdductorMuscle4 | SRR33418589 | 9419443 | *O. bimaculoides* | Tentacle_Ctrl2 | SRR33418568 | 50283992 |
| *C. virginica* | AdductorMuscle9 | SRR33418588 | 9807973 | *O. bimaculoides* | Tentacle_Ctrl3 | SRR33418567 | 36235581 |
| *C. virginica* | AdductorMuscle10 | SRR33418587 | 9593441 | *O. bimaculoides* | Tentacle_Reg1 | SRR33418566 | 21210311 |
| *C. virginica* | AdductorMuscle11 | SRR33418586 | 9923013 | *O. bimaculoides* | Tentacle_Reg2 | SRR33418565 | 39412129 |
| *C. virginica* | AdductorMuscle12 | SRR33418585 | 9571981 | *O. bimaculoides* | Tentacle_Reg3 | SRR33418564 | 41426978 |
| *C. virginica* | AdductorMuscle13 | SRR33418584 | 9732943 | *D. melanogaster* | Ovary_WT1 | SRR11680898 | 39157763 |
| *C. virginica* | AdductorMuscle14 | SRR33418583 | 9476406 | *D. melanogaster* | Ovary_WT2 | SRR11680899 | 31733032 |
| *C. virginica* | AdductorMuscle15 | SRR33418582 | 9482977 | *D. melanogaster* | Ovary_WT3 | SRR11680900 | 34563946 |
| *C. virginica* | AdductorMuscle16 | SRR33418580 | 9271958 | *D. melanogaster* | Testes_WT1 | SRR12313524 | 19163553 |
| *C. virginica* | Gill1 | SRR33418579 | 14050252 | *D. melanogaster* | Testes_WT2 | SRR12313525 | 16768121 |
| *C. virginica* | Gill2 | SRR33418578 | 13712068 | *D. melanogaster* | Testes_WT3 | SRR12313526 | 18081329 |
| *C. virginica* | Gill3 | SRR33418577 | 14609013 | *M. musculus* | Ovary_Ctrl1 | SRR13848060 | 20989881 |
| *C. virginica* | Gill4 | SRR33418576 | 14100857 | *M. musculus* | Ovary_Ctrl3 | SRR13848061 | 43772792 |
| *C. virginica* | Gonad1 | SRR33418575 | 13244351 | *M. musculus* | Ovary_Ctrl6 | SRR13848062 | 26697670 |
| *C. virginica* | Gonad2 | SRR33418574 | 12940268 | *M. musculus* | Testis1 | SRR24821465 | 7159893 |
| *C. virginica* | Gonad3 | SRR33418573 | 13617227 | *M. musculus* | Testis2 | SRR24821466 | 9348548 |
| *C. virginica* | Gonad4 | SRR33418572 | 13226919 | *M. musculus* | Testis3 | SRR24821469 | 9546916 |
| *C. virginica* | CvirMantleSnRNA_1 | SRR33450334 | 41392647 | *M. musculus* | Testis4 | SRR24821470 | 10192831 |
| *C. virginica* | CvirMantleSnRNA_2 | SRR33450334 | 41392647 | *S. mediterranea* | Salami_Section10 | SRR12426750 | 27373271 |
| *R. venosa* | Pre-Competent_Larvae1 | SRR5931826 | 10157033 | *S. mediterranea* | Salami_Section9 | SRR12426751 | 7597498 |
| *R. venosa* | Pre-Competent_Larvae2 | SRR5931827 | 10827765 | *S. mediterranea* | Salami_Section8 | SRR12426752 | 12112061 |
| *R. venosa* | Pre-Competent_Larvae3 | SRR5931828 | 10601339 | *S. mediterranea* | Salami_Section7 | SRR12426753 | 9028748 |
| *R. venosa* | Competent_Larvae1 | SRR5931829 | 12156108 | *S. mediterranea* | Salami_Section6 | SRR12426754 | 6048316 |
| *R. venosa* | Competent_Larvae2 | SRR5931830 | 11451480 | *S. mediterranea* | Salami_Section5 | SRR12426755 | 8503488 |
| *R. venosa* | Competent_Larvae3 | SRR5931831 | 10023123 | *S. mediterranea* | Salami_Section4 | SRR12426756 | 6511945 |
| *R. venosa* | Post-Larvae1 | SRR5931832 | 10678504 | *S. mediterranea* | Salami_Section3 | SRR12426757 | 11365797 |
| *R. venosa* | Post-Larvae2 | SRR5931833 | 11622298 | *S. mediterranea* | Pharynx_Section12 | SRR12426758 | 8309356 |
| *R. venosa* | Post-Larvae3 | SRR5931834 | 12087773 | *S. mediterranea* | Salami_Section11 | SRR12426759 | 8187802 |
| *M. yessoensis* | SmoothMuscle1 | SRR18577892 | 10486504 | *S. mediterranea* | Salami_Section2 | SRR12426760 | 8476769 |
| *M. yessoensis* | SmoothMuscle2 | SRR18577891 | 11914196 | *S. mediterranea* | Salami_Section1 | SRR12426761 | 5886612 |
| *M. yessoensis* | SmoothMuscle3 | SRR18577890 | 10417121 | *L. stagnalis* | Control1 | SRR12675210 | 27713142 |
| *M. yessoensis* | StriatedMuscle1 | SRR18577889 | 12011743 | *L. stagnalis* | Control2 | SRR12675206 | 25417785 |
| *M. yessoensis* | StriatedMuscle2 | SRR18577888 | 14665041 | *L. stagnalis* | Control3 | SRR12675247 | 31242967 |
| *M. yessoensis* | StriatedMuscle3 | SRR18577887 | 11196753 | *L. stagnalis* | Control4 | SRR12675229 | 19960951 |
| *M. gigas* | AdductorMuscle1 | SRR6489635 | 27407026 | *L. stagnalis* | Wounded1 | SRR12675230 | 27023963 |
| *M. gigas* | AdductorMuscle2 | SRR6489636 | 29698653 | *L. stagnalis* | Wounded2 | SRR12675245 | 23379402 |
| *M. gigas* | Gonad1 | SRR6489633 | 28031758 | *L. stagnalis* | Wounded3 | SRR12675236 | 19302665 |
| *M. gigas* | Gonad2 | SRR6489634 | 30366538 | *L. stagnalis* | Wounded4 | SRR12675227 | 14728302 |
| *H. rufescens* | AdductorMuscle | SRR33418563 | 51801246 |  |  |  |  |

**Supplemental Table 3:**
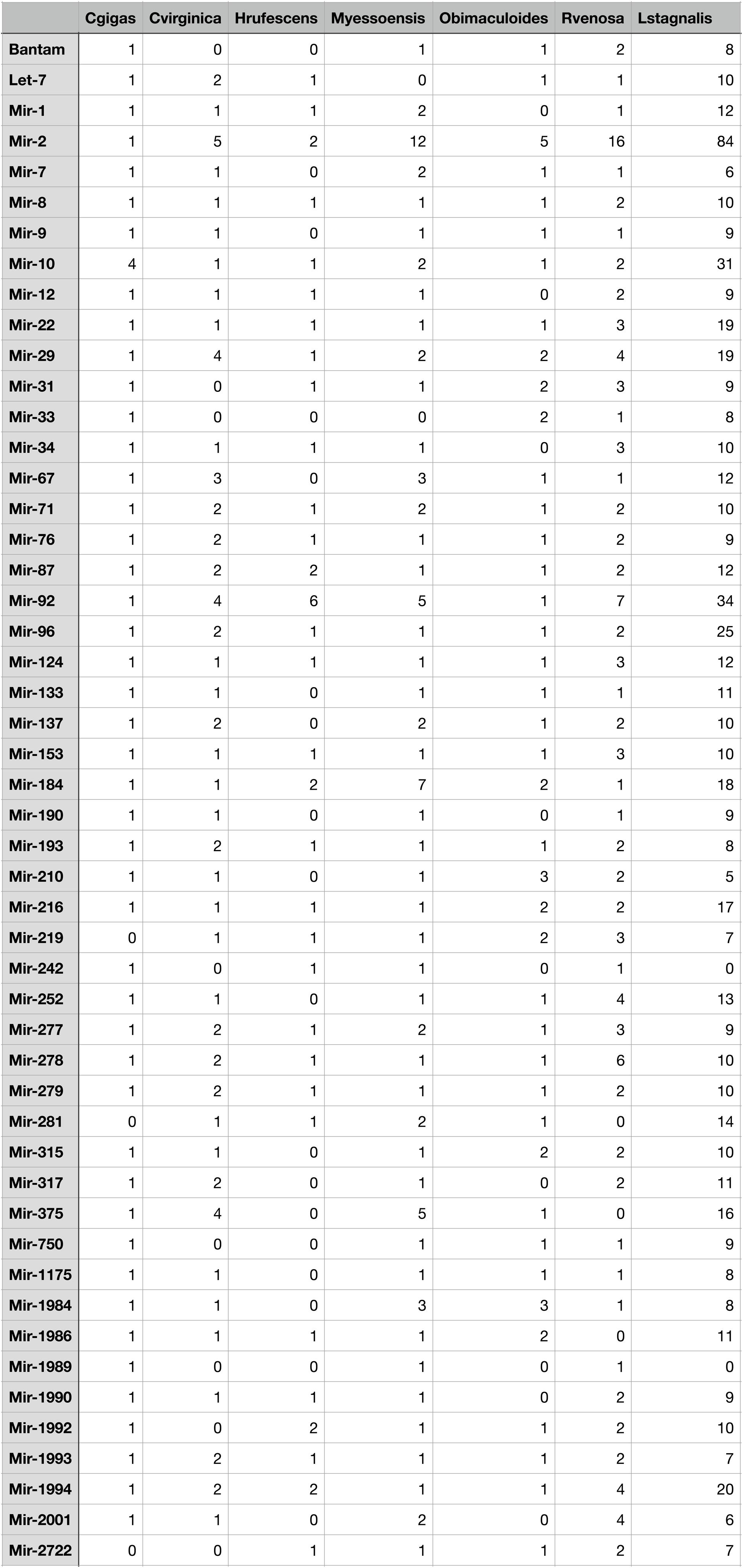
Overview of miRNA Families in Mollusca. The top 50 conserved miRNA sequences identified amongst the mollusks examined.

|  | Cgigas | Cvirginica | Hrufescens | Myessoensis | Obimaculoides | Rvenosa | Lstagnalis |
| --- | --- | --- | --- | --- | --- | --- | --- |
| Bantam | 1 | 0 | 0 | 1 | 1 | 2 | 8 |
| Let-7 | 1 | 2 | 1 | 0 | 1 | 1 | 10 |
| Mir-1 | 1 | 1 | 1 | 2 | 0 | 1 | 12 |
| Mir-2 | 1 | 5 | 2 | 12 | 5 | 16 | 84 |
| Mir-7 | 1 | 1 | 0 | 2 | 1 | 1 | 6 |
| Mir-8 | 1 | 1 | 1 | 1 | 1 | 2 | 10 |
| Mir-9 | 1 | 1 | 0 | 1 | 1 | 1 | 9 |
| Mir-10 | 4 | 1 | 1 | 2 | 1 | 2 | 31 |
| Mir-12 | 1 | 1 | 1 | 1 | 0 | 2 | 9 |
| Mir-22 | 1 | 1 | 1 | 1 | 1 | 3 | 19 |
| Mir-29 | 1 | 4 | 1 | 2 | 2 | 4 | 19 |
| Mir-31 | 1 | 0 | 1 | 1 | 2 | 3 | 9 |
| Mir-33 | 1 | 0 | 0 | 0 | 2 | 1 | 8 |
| Mir-34 | 1 | 1 | 1 | 1 | 0 | 3 | 10 |
| Mir-67 | 1 | 3 | 0 | 3 | 1 | 1 | 12 |
| Mir-71 | 1 | 2 | 1 | 2 | 1 | 2 | 10 |
| Mir-76 | 1 | 2 | 1 | 1 | 1 | 2 | 9 |
| Mir-87 | 1 | 2 | 2 | 1 | 1 | 2 | 12 |
| Mir-92 | 1 | 4 | 6 | 5 | 1 | 7 | 34 |
| Mir-96 | 1 | 2 | 1 | 1 | 1 | 2 | 25 |
| Mir-124 | 1 | 1 | 1 | 1 | 1 | 3 | 12 |
| Mir-133 | 1 | 1 | 0 | 1 | 1 | 1 | 11 |
| Mir-137 | 1 | 2 | 0 | 2 | 1 | 2 | 10 |
| Mir-153 | 1 | 1 | 1 | 1 | 1 | 3 | 10 |
| Mir-184 | 1 | 1 | 2 | 7 | 2 | 1 | 18 |
| Mir-190 | 1 | 1 | 0 | 1 | 0 | 1 | 9 |
| Mir-193 | 1 | 2 | 1 | 1 | 1 | 2 | 8 |
| Mir-210 | 1 | 1 | 0 | 1 | 3 | 2 | 5 |
| Mir-216 | 1 | 1 | 1 | 1 | 2 | 2 | 17 |
| Mir-219 | 0 | 1 | 1 | 1 | 2 | 3 | 7 |
| Mir-242 | 1 | 0 | 1 | 1 | 0 | 1 | 0 |
| Mir-252 | 1 | 1 | 0 | 1 | 1 | 4 | 13 |
| Mir-277 | 1 | 2 | 1 | 2 | 1 | 3 | 9 |
| Mir-278 | 1 | 2 | 1 | 1 | 1 | 6 | 10 |
| Mir-279 | 1 | 2 | 1 | 1 | 1 | 2 | 10 |
| Mir-281 | 0 | 1 | 1 | 2 | 1 | 0 | 14 |
| Mir-315 | 1 | 1 | 0 | 1 | 2 | 2 | 10 |
| Mir-317 | 1 | 2 | 0 | 1 | 0 | 2 | 11 |
| Mir-375 | 1 | 4 | 0 | 5 | 1 | 0 | 16 |
| Mir-750 | 1 | 0 | 0 | 1 | 1 | 1 | 9 |
| Mir-1175 | 1 | 1 | 0 | 1 | 1 | 1 | 8 |
| Mir-1984 | 1 | 1 | 0 | 3 | 3 | 1 | 8 |
| Mir-1986 | 1 | 1 | 1 | 1 | 2 | 0 | 11 |
| Mir-1989 | 1 | 0 | 0 | 1 | 0 | 1 | 0 |
| Mir-1990 | 1 | 1 | 1 | 1 | 0 | 2 | 9 |
| Mir-1992 | 1 | 0 | 2 | 1 | 1 | 2 | 10 |
| Mir-1993 | 1 | 2 | 1 | 1 | 1 | 2 | 7 |
| Mir-1994 | 1 | 2 | 2 | 1 | 1 | 4 | 20 |
| Mir-2001 | 1 | 1 | 0 | 2 | 0 | 4 | 6 |
| Mir-2722 | 0 | 0 | 1 | 1 | 1 | 2 | 7 |

